# CRISPR-assisted targeted enrichment-sequencing (CATE-seq)

**DOI:** 10.1101/672816

**Authors:** Xinhui Xu, Qiang Xia, Shuyan Zhang, Jinliang Gao, Wei Dai, Jian Wu, Jinke Wang

## Abstract

The current targeted sequencing of genome is mainly dependent on various hybridization-based methods. However, the hybridization-based methods are still limited by the intrinsic shortcomings of nucleic acid hybridization. This study developed a new CRISPR-based targeted sequencing technique, CRISPR-assisted targeted enrichment-sequencing (CATE-seq). In this technique, the input genomic DNA (gDNA) was firstly bound by a complex of dCas9 and capture sgRNA (csgRNA). The DNA-dCas9-csgRNA complex was then captured on magnetic beads through an easy room-temperature annealing between a short universal capture sequence (24 bp) at the 3′ end of csgRNA and capture oligonucleotide coupled on magnetic beads. The enriched DNAs were finally analyzed by next generation sequencing. Using this technique, three different scales of targeted enrichments were successfully performed, including enriching 35 target exons of 6 genes from 6 gDNA samples with 54 csgRNAs, 339 target exons of 186 genes from 9 gDNA samples with 367 csgRNAs, and 2031 target exons of 451 genes from 2 gDNA samples with 2302 csgRNAs. This technique has several significant advantages over the current hybridization-based methods, including high simplicity, specificity, sensitivity, throughput, and scalability.

## INTRODUCTION

The present precision medicine is increasingly dependent on DNA information. More and more clinical DNA samples were analyzed for finding the diagnostic and therapeutic markers of diseases. Especially, analysis of cell-free DNA provides new chance to disease diagnosis by liquid biopsy. DNA analysis can find genetic and epigenetic alterations in genomes, such as single-nucleotide variants (SNVs), copy-number variants (CNVs), translocations, and methylation. Along with the advance of DNA sequencing techniques, next-generation sequencing (NGS) provides a powerful tool for decoding DNA information at the genome scale and single-base resolution. The NGS-based DNA analysis can provide some valuable information. First, the whole genome sequencing (WGS) can be used to systematically identify cancer-related polymorphism and signature mutations. Second, the targeted gene panel sequencing (TGPS) can be used to detect the known pathogenic mutations for diagnosis and prognosis. Third, the hydrogen sulfite sequencing can be used to find the alterations of important epigenetic marker of DNA methylation. Finally, WGS can also be used to characterize another important epigenetic marker, chromatin openness state (ATAC-seq), which is determined by epigenetic modifications of nucleosomes such as methylation and acetylation. The genetic alterations occurred in exons may produce the dysfunctional proteins or RNAs. The genetic and epigenetic alterations occurred in regulatory regions may lead to dysregulation of gene expression. Therefore, all these alterations disclosed by DNA sequencing may contribute to the occurrence of diseases.

Although WGS provides more complete genomic DNA (gDNA) information, it is low cost-effectiveness. The targeted DNA sequencing (TDS) is therefore increasingly adopted such as whole exome sequencing (WES) (1) and whole regulome sequencing (WRS) (2). TDS has several advantages over WGS. First, TDS is more cost-effective. Second, TDS facilitates higher sample throughput than WGS, allowing to reveal biologically important sequence variation in a large number of individuals (3). Third, TDS improves accuracy by optimizing the read depth coverage and reducing the complexity of DNA to be sequenced. These advantages make TDS applicable to clinics. For instance, in the CAncer Personalized Profiling by deep Sequencing (CAPP-Seq) (4,5), the recurrent mutations in a particular cancer type that were collected in the Catalogue of Somatic Mutations in Cancer (COSMIC) and The Cancer Genome Atlas (TCGA) could be identified by hybridizing the tumor biopsy DNA with the selector probe set and deep sequencing. CAPP-Seq was thus applied to monitoring tumor burden, prognostic indicator, and biopsy-free tumor genotyping (5). TDS is therefore more valuable to the current biomedicine.

Currently, the targeted enrichment of genomic DNA (gDNA) can be realized in four approaches (6–9). The first is DNA hybridization on solid surface such as DNA chip or microarray (10–15). The second is DNA hybridization in solution with biotin-labeled capture probes (DNA or RNA) and the hybrids were isolated with streptavidin-coupled magnetic beads (4,5,16,17). The third is the targeted PCR amplification (8,18,19). The fourth is the DNA capture via molecular inversion probe circularization (20,21). The first two approaches have high throughput due to using large numbers of capture probes. The targeted PCR amplification has low throughput due to the limitation of multiplexed PCR and is therefore often used to detect limited numbers of disease-causing sites or genes. Currently, the in-solution hybridization becomes the main approach of targeted sequencing due to free of expensive chips/microarrays and the corresponding additional equipment. The in-solution hybridization approach has been therefore widely adopted by most commercialized WES kits (e.g. Agilent’s SureSelect Human All Exon, Roche/Nimblegen’s SeqCap EZ Exome Library, Illumina’s TruSeq Exome Enrichment, and iGeneTech’s AIwholeExome). However, in-solution hybridization approach is still challenged by the intrinsic limitations of nucleic acid hybridization, such as DNA denaturing and hybridizing for a long time at high temperature, unavoidable nonspecific hybridization that produces high background signal and lowers enrichment specificity, and consumption of large numbers of single-stranded biotinylated capture oligonucleotides that have to be repeatedly chemically synthesized. Therefore, new hybridization-free and cost-effective targeted enrichment sequencing techniques are still in demand.

CRISPR is originally an immune system of bacteria to destroy the invaded microphage DNA by enzymatically digestion. The system has already been developed into a high-efficiency gene editing tool (22,23). In addition, the system has also been developed as a new kind of gene regulation tool. For example, the deactivated or “dead” Cas9 (dCas9) and its associated single guide RNA (sgRNA) have been most widely used to regulate gene expression in recent years (24–30). In these applications of CRISPR/Cas9 technique, both dCas9 and sgRNA were widely engineered. Compared with Cas9 engineering, sgRNA is more simple, flexible, and efficient to redesign. SgRNA was engineered by extending new various functional sequences at its 3’ end for binding various RNA-binding proteins, such as MS2 bacteriophage coat proteins (31,32), Pumilio/FBF (PUF) (33), MCP, PCP, and Com (34). These RNA-binding proteins were fused with transcription-activation domains such as VP64 (33), VP64-HSF1(31,32), and p65-HSF1 (33). SgRNA was also engineered to include modified riboswitches for recognizing specific signals such as drug (35). The chimeric sgRNAs greatly widened the application of CRISPR technique, suggesting that chimeric sgRNAs has great potential in exploration of novel applications of CRISPR technique.

This study developed a new CRISPR-based targeted enrichment technique. The 3′ end of sgRNA was re-designed to contain a universal short capture sequence (CS). This kind of special sgRNA was named as capture sgRNA (csgRNA). A corresponding constant biotinylated single-stranded capture oligonucleotide (CP) was anchored on the surface of streptavidin-coated magnetic beads. In this technique, input dsDNA library was firstly bound by the dCas9-csgRNA complexes, and the formed dsDNA-dCas9-csgRNA complexes were then isolated with the CP-coupled magnetic beads via an easy room-temperature annealing between CS and CP. The enriched DNA was finally analyzed by NGS. This technique was thus named as the CRISPR-assisted targeted enrichment-sequencing (CATE-seq). This technique was fully verified by three different scales of targeted enrichments.

## MATERIALS AND METHOD

### Cell culture

All cells were purchased from the Cell Culture Bank of the Chinese Academy of Sciences (Shanghai, China). Cells were cultivated with the DMEM (HyClone) (293T, HepG2, HeLa, SiHa, and C-33A) or RPMI-1640 (HyClone) (HL7702) medium supplemented with 10% fetal bovine serum (HyClone), 100 U/mL penicillin, and 100 μg/mL streptomycin (Biosharp). Cells were incubated at 37°C in a 5% CO_2_ incubator.

### DNA extraction

Cells were washed twice with PBS and collected by trypsinization. The gDNA was extracted from cells using the TIANamp Genomic DNA Kit (Tiangen). The extracted gDNA was quantified with spectrometry and stored at −80°C until use.

### DNA tagmentation

The barcoded transposons were prepared by annealing two complementary oligonucleotides, Barcode 1–7 and ME oligo (Table S1). Seven barcoded transposons (ME-B-1–7) (Table S2) were thus prepared. The Tn5 transposome was then assembled in a 20-μL reaction containing 1 μM transposon, 1× TPS buffer (Robustnique, China), and 5 U of Robust Tn5 transposase (Robustnique, China). The reaction was incubated at 25°C for 30 min. The assembled Tn5 transposomes (T-B-1–7) were stored at −20°C until use. The 293T, HepG2, HL7702, C-33a, SiHa, and HeLa gDNAs were tagmented with the Tn5 transposomes T-B-1 to T-B-6, respectively. The 293T gDNA was also tagmented with the Tn5 transposome T-B-7 (293Tm). A 30-μL tagmentation reaction contained 200 ng gDNA, 5 μL transposome, and 1× LM buffer (Robustnique, China). The reaction was incubated at 55°C for 15 min. The reaction contained no transposome was used as control. The reactions were run with agarose gel. The fragments of 200-1000 bp were recovered from gel and used as the input gDNA of targeted enrichment.

### SgRNA preparation

The sequences of target transcripts were obtained from hg19 human genome with the UCSC browser. The non-redundant full-length CDS sequences of target genes were obtained with a home-made script. The 20-bp flanking genomic sequences were added to both ends of the CDSs shorter than 70 bp. SgRNAs were then designed for all CDSs using the CHOPCHOP program (Table S3). CsgRNAs were prepared with *in vitro* transcription. A three-round fusion PCR amplification was used to prepare DNA template of csgRNA transcription (see details as supplementary method). PCR primers were shown in Table S4. The final PCR product was purified by gel recovery. A 20-μL transcription reaction contained 50 U T7 RNA polymerase (New England Biolabs, NEB), 1× T7 RNA polymerase buffer (NEB), 4 mM each rNTP (NEB), and 500 ng DNA template. The reaction was incubated at 37°C overnight. RNA was purified with the Trizol method and dissolved in RNase-free ddH_2_O. CsgRNA was diluted to a working concentration of 15 ng/μL and stored at −80°C until use.

### Targeted enrichment

The dCas9 protein (NEB) was diluted to 0.3 μM with stock solution and stored at −20°C. CsgRNA pools were prepared by mixing various csgRNAs No. 1–11, 12–25, 26–40, and 41–54, respectively. Each csgRNA pool contained about 7.5 ng of each csgRNA. Five microliters of Dynabead-M-280-streptavidin (Invitrogen) were washed three times with 50 μL of PBS-BSA solution (PBS solution with 0.5% BSA) and resuspended in 50 μL of 1×dCas9 buffer (NEB) with 0.4 μM CP (5′-Biotin-TTTTT TTGCA TCTGG TATTC GTAAG GTTCC G-3′). The beads were rotated for 1 h at room temperature (RT) and then washed three times with 50 μL of 1×dCas9 buffer. Each csgRNA pool was added with 4 μL of 0.3 μM dCas9, 4 U RNase Inhibitor (ThermoFisher), and 2.5 μL of 10× dCas9 Buffer, and supplemented to a total of 25 μL with DEPC-treated ddH_2_O. The mixture was incubated at 25°C for 10 min to form dCas9-csgRNA complexes. Five microliters of input gDNA (about 5 ng) was then directly added into above dCas9-csgRNA mixture and incubated at 37°C for 30 min. The gDNA-dCas9-csgRNA mixture was then added to the above prepared CP-coupled magnetic beads and rotated at RT for 1 h. Subsequently, the beads were washed three times with 50 μL of PBS-BSA solution. Finally, the beads were resuspended in 30 μL of TE buffer (pH8.1) and incubated at 85°C for 5 min. The beads were then rapidly placed on a magnet and the supernatant was quickly transferred into a clean EP tube as the enriched target DNA. The enriched gDNA was stored at −20°C for later use.

### DNA sequencing

The NGS library was constructed using a SALP method developed by our lab (36). A single-strand adaptor (SSA) was prepared by annealing two oligonucleotides, SA and SA-3N (Table S5). Ten microliters of enriched gDNA were denatured at 95°C for 5 min and immediately chilled on ice. A 20-μL ligation reaction contained 10 μL denatured enriched gDNA, 1 U T4 DNA ligase (Invitrogen), 1× ligation buffer, and 0.5 μM SSA. The reaction was incubated overnight at 16°C. A 40-μL extension reaction contained 20 μL ligation reaction and 1× rTaq mix (Takara). The reaction was incubated at 72° C for 15 min. A 50-μL PCR reaction contained 20 μL extension reaction, 1× NEB Next® Q5® Hot Start HiFi PCR Master Mix (NEB, M0543S), 0.2 μM Universal Primer (Table S4), and 0.2 μM IP15 (Table S4). The PCR program was as follows: 98°C 5 min, 17 cycles of 98°C 10 s, 65°C 30 s and 72°C 1 min, and 72°C 5 min. The PCR product was run with 1.5% agarose gel. The DNA smear of 200–1000 bp was recovered with the Axygen DNA Gel Recovery Kit (Axygen). The recovered DNA was quantified using Qubit. A total of 7 DNA libraries were mixed in the same quality (ng) and sequenced in a lane of Hiseq-4000, using double-ended 150-bp sequencing.

### Reads analysis

The reads in the raw CATE-seq data were assigned to DNA samples according to correct barcode sequence using a home-made Python script. The reads numbers of each DNA sample were counted. The following parameters were used in assigning the raw CATE-seq data to form the fastq files for each DNA sample: (1) At most 5 non-contiguous indefinite bases (N) were allowed; (2) At most one non-terminal N base was allowed; (3) Barcode sequence consisted of 6 nucleotides. The assigned reads data in fastq files were then mapped to the reference human genome (hg19) with the bowtie (Centos5.5 operating system). The resulted sam files were converted to bam files with samtools. The resulted bam files were then sorted with samtools. Finally, the sorted bam files were converted to BigWig files with the bedGraphToBigWig for visualizing the mapping results with the UCSC genome browser.

### TERT enrichment

A 235-bp wild-type TERT promoter DNA fragment (TERT-P) and a mutant TERT promoter DNA fragment with a C/T mutation at position −158 (TERT-P-mut) were prepared with PCR amplification using primers of TERT-PF, TERT-PR, and TERT-Mut-R (Table S6) (see details as supplementary methods). A sgRNA targeting the mutant TERT promoter was designed by using the GGG at position from −162 bp to −164 bp as PAM. The sequence of sgRNA target and PAM are 5′-TCCCC GGCCC AGCCC C**T**TCC GGG (from −142 to −164; T in bold is the mutant base). The csgRNA was then prepared by *in vitro* transcription using the template produced by three-round fusion PCR. The PCR primers were F1, R, sgR1, TERT-sgRNA-F2, and TERT-sgRNA-F3 (Table S4 and S6).

A cocktail of wild-type and mutant TERT promoter DNA fragments was prepared by mixing the same quality of TERT-P and TERT-P-mut (0.2 ng/μL). The TERT-P-mut was then enriched from the cocktail with CATE. The enriched DNA was amplified with rTaq mix (Takara) for 20 cycles using primers TERT-PF and TERT-PR. The PCR products were purified with a PCR clean kit and ligated into a T vector. Twenty positive colonies were sequenced. The ratio of the mutant clones to the wild-type clones was calculated.

In another enrichment, the TERT-P and TERT-P-mut were mixed at the various ratios (1:1, 1:10, 1:100, 1:1,000, 1:10,000, 1:100,000, and 1:1,000,000). The total DNA concentration of all mixtures was 0.2 ng/μL. Each mixture was enriched with the csgRNA targeting TERT-P-mut. The enriched DNA was analyzed with ARMS-qPCR (37,38) using designed extended primers (Table S6) (see details as supplementary methods). The ratio of mutants in the enriched DNA was determined according to the ARMS-qPCR results. The percentage and enrichment fold of mutant TERT promoter in the enriched DNA was calculated according to the Ct values.

## RESULTS

### Principle of CRISPR-assisted targeted enrichment

The principle of CRISPR-assisted targeted enrichment (CATE) is schematically illustrated in Figure 1. The tagmented gDNA were used as the input DNA of CATE (Figure 1A). In this method, a capture sgRNA (csgRNA) was developed by extending a short CS (5′-CGGAA CCTTA CGAAT ACCAG ATGC-3′) at the 3′ end of normal sgRNA (Figure 1B). Correspondingly, a complementary oligonucleotide was used as CP and coupled on magnetic beads via biotin-streptavidin interaction (Figure 1B). SgRNAs targeting DNAs of interest were designed and csgRNA were prepared by *in vitro* transcription. The input DNA was mixed with the pre-assembled dCas9-csgRNA complex, allowing the dCas9-csgRNA complex to bind its targets. The mixture was then incubated with the CP-coupled magnetic beads and the DNA-dCas9-csgRNA complex was magnetically isolated. The captured DNA was analyzed with our recently developed SALP-seq method (Figure 1A).

**Figure 1.**
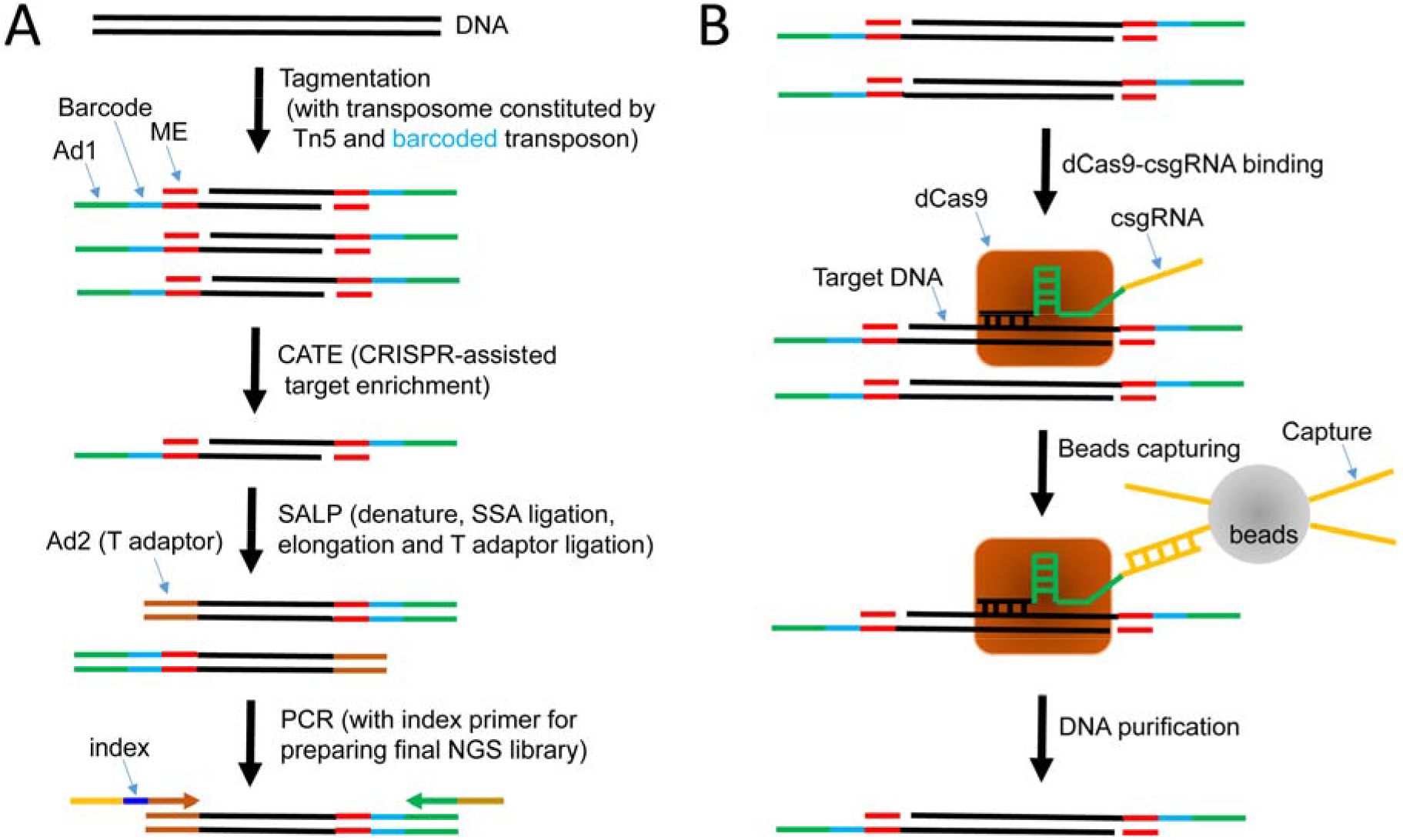
Principle of CRISPR-assisted targeting enrichment (CATE). (A) Schematic show of procedures of CATE-seq. SSA, single strand adaptor; SALP, single-stranded adaptor library preparation. ME, mosaic end; Ad1, adaptor 1 (Universal Primer annealing site); Ad2, adaptor 2 (Index Primer annealing site); NGS, next-generation sequencing. (B) Schematic show of principle of CATE. dCas9, deactivated Cas9; csgRNA, capture sgRNA. Capture, capture oligo immobilized on the surface of magnetic beads, which can be annealed with the capture sequence at the 3′ end of csgRNA.

### DNA tagmentation, CATE and NGS library construction

The gDNAs of six cell lines (293T, HepG2, HL7702, HeLa, SiHa, and C-33A) were tagmented with different barcoded transposons (Table S2). The 293T gDNA was also tagmented by another barcode transposon to form an input DNA named as 293Tm. The agarose gel electrophoresis revealed that all gDNA samples were successfully tagmented into gDNA fragments (Figure S1A). The DNA fragments of 200-1000 bp were then isolated by gel recovery (Figure S1A) and enriched with CATE. The enriched DNA was used to constructed NGS library using SALP method (Figure 1A). The detection of final PCR products with agarose gel electrophoresis showed that all enriched DNA was successfully amplified (Figure S1B). The DNA fragments of 200–1000 bp were then recovered and analyzed with NGS. Two negative controls, a CATE without csgRNA and a CATE with sonicated gDNA, did not produce PCR products.

### Cloning sequencing of NGS library

In fact, the feasibility of CATE protocol was firstly testified with the 293T gDNA. The prepared NGS library was detected by cloning sequencing. Fifty colony PCR-identified positive clones were randomly selected to sequence (Figure S2). The sequences (see supplementary sequences) were mapped to the reference genome (hg19). The results indicated that all clones could be mapped to the sgRNA-targeting CDSs of 6 genes (Figure S3 and S4). All clones contained the sgRNA targets (see supplementary sequences). In addition, the number of clones of a gene was proportional to the number of sgRNAs designed for the gene. The CDSs with two sgRNAs were preferentially sequenced in the cloning sequencing (31 clones). These results demonstrated the feasibility, reliability, and specificity of CATE method. The gDNA of other five cell lines were then subjected to CATE following the established protocol.

### NGS and data analysis

Seven NGS libraries of 293T, HepG2, HL7702, HeLa, SiHa, C-33A, and 293Tm were mixed together at the same micrograms to form a pooled final NGS library. This library was then sequenced with a lane of Hiseq-4000. As a result, a total of 163,270,664 reads were obtained, in which 135,607,186 reads had correct barcode (83% of total reads) (Table S7). After mapping, 124,699,943 reads were mappable (76% of total reads and 92% of correct barcode reads) (Table S7). After assigning all reads to DNA samples, the total reads, mappable reads, and targeted reads of each sample were counted (Table 1; Figure 2A). The targeted reads were defined as the reads of the pair-end sequenced DNA fragments that contained the sgRNA targets. The results indicated that as many as 112,760,368 reads were targeted reads (92% of 124,699,943 mappable reads) (Table S7). These data indicate that CATE has high target specificity. The DNA fragments with all mappable pair-end reads had the similar length distribution (Figure S5). The similar reads distribution, mappability and targeting among 7 samples (Table 1) reveal that the CATE method has high reproducibility.

**Table 1.** Assignment, mappability and targeting of CATE-seq reads among DNA samples.

| DNA sample | Barcode number | Barcode sequence | Index | Total Reads | Mappable reads | % of mappable reads | Targeted reads | % of targeted reads |
| --- | --- | --- | --- | --- | --- | --- | --- | --- |
| 293T | Barcode 1 | ATCACG | TGACAT | 20140283 | 18723387 | 92.96% | 16579325 | 88.55% |
| HepG2 | Barcode 2 | CGATGT | TGACAT | 19973759 | 18509682 | 92.67% | 17219144 | 93.03% |
| HL7702 | Barcode 3 | TGACCA | TGACAT | 19118758 | 17826330 | 93.24% | 15846385 | 88.89% |
| C-33a | Barcode 4 | CAGATC | TGACAT | 18346510 | 17122798 | 93.33% | 15677681 | 91.56% |
| SiHa | Barcode 5 | GATCAG | TGACAT | 20417029 | 18528454 | 90.75% | 16683647 | 90.04% |
| HeLa | Barcode 6 | CTTGTA | TGACAT | 18600942 | 16619942 | 89.35% | 15136726 | 91.08% |
| 293Tm | Barcode 7 | GGCTAC | TGACAT | 19009905 | 17369350 | 91.37% | 15617460 | 89.91% |

**Figure 2.**
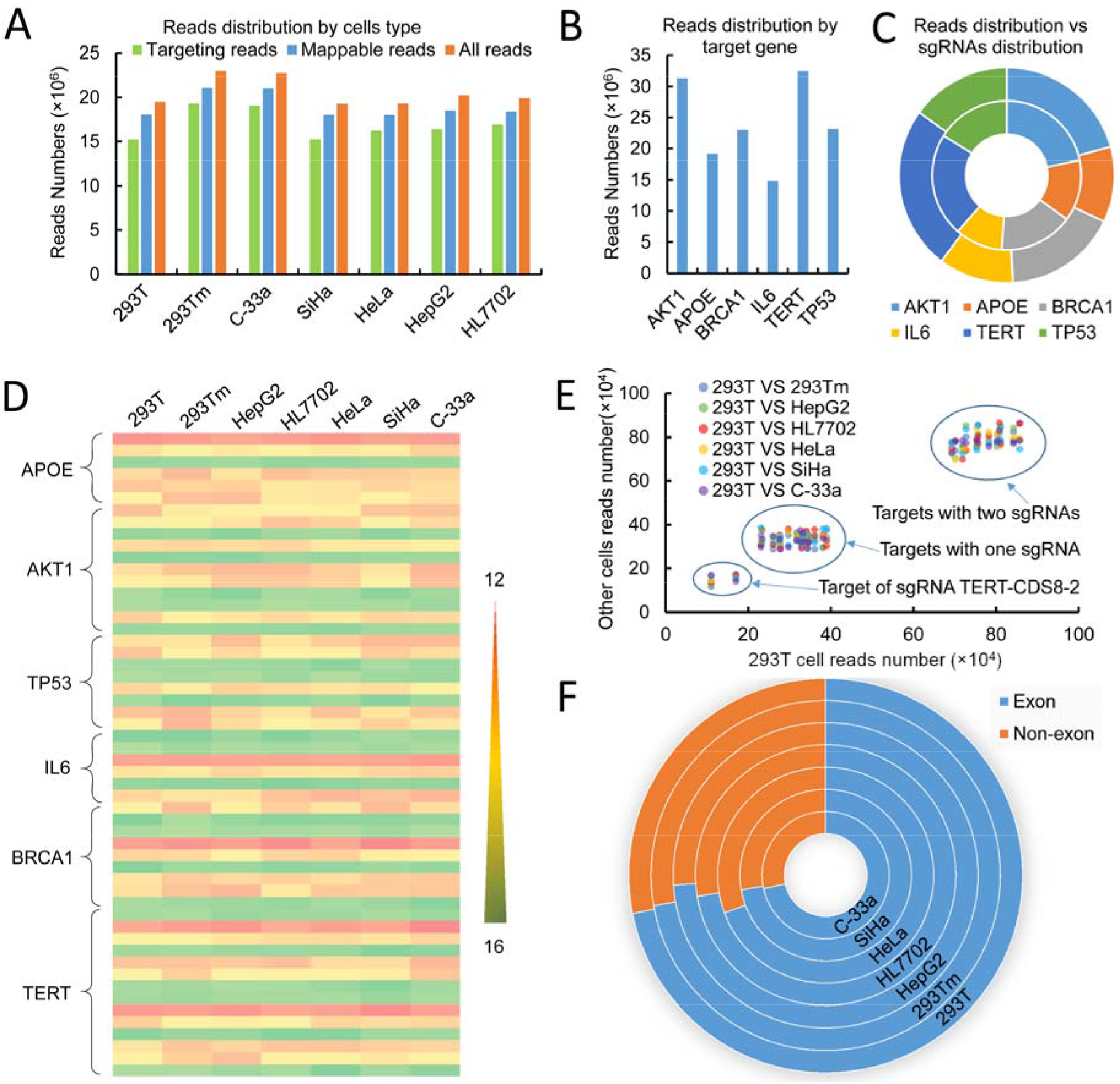
Statistical results of CATE-seq reads. (A) Distribution of all, mappable, and targeted reads in 7 DNA samples. (B) Distribution of all mappable reads in six genes. (C) Reads distribution (%) versus sgRNA distribution (%) in six genes. Outer, sgRNA distribution; inner, reads distribution. (D) Reads distribution in all sgRNA targets in 6 genes in 7 DNA samples. The log2 value of reads number was used. Each raw represents one sgRNA target. (E) Comparison of reads of 293T DNA sample with those of other DNA samples. The dots with the lowest reads number refers to the reads enriched by sgRNA TERT-CDS8-2. (F) Distribution of all bases covered by targeting reads in exon and non-exon.

### Characterization of reads assignment to target genes

The number of reads assigned to 6 target genes was then counted (Figure 2B). The results indicated that the number of reads assigned to a particular gene was correlated with the numbers of sgRNAs designed for the gene (Figure 2C), showing the enrichment effect of CATE. The reads mapped to all sgRNA-targeting CDSs of 6 genes in all DNA samples were then counted. The results demonstrated that all sgRNAs effectively enriched their targets, all sgRNAs had the stable enrichment efficiency to their targets in all DNA samples, and the enrichment fold of a CDS was related with the number of its targeting sgRNAs (Figure 2D). The reads distribution was also checked by counting the reads of each target (Table S8). Based on this data, the enrichment efficiency of all sgRNA in all DNA samples were further evaluated by comparing the reads of 293T DNA sample with those of other DNA samples. The results indicate that only sgRNA TERT-CDS8-2 showed a similar lowest enrichment efficiency in 7 samples (Figure 2E). All other sgRNAs had the similar enrichment efficiency in all samples (Figure 2E). The targets targeted by two closely neighbored sgRNAs were enriched twice over those targeted by one sgRNA (Figure 2E). Finally, all bases covered by targeted reads in exon and non-exon were counted and compared. It was found that the bases in exons and non-exons occupied 75% and 25% in all 7 samples, respectively (Figure 2F).

### Specificity of reads targeting

The global reads distribution was displayed with the CIRCOS Circular Genome Data Visualization (Figure 3). It revealed that most of reads located at the sgRNA-targeted loci. However, there were still some “off-target” reads (11,939,575 reads; Table S7) distributing in other genomic regions (Figure 3). Nevertheless, these “off-target” reads were evenly distributed in the whole genome with very low reads density. Further comparison revealed that the distribution of these “off-target” reads in genome had no relationship with the predicted off-targets (Figure 3). These “off-target” reads may come from the very low amount of non-specific absorption of gDNA to beads. These data also revealed the high specificity of CATE. The sgRNA and reads distribution in target genes were also displayed by the UCSC Genome Browser (39) (Figure 4). It was found that reads were highly enriched at the loci of five target genes in all DNA samples. More importantly, reads were highly enriched in the sgRNAs-targeted exons. The exons targeted by two sgRNAs were more highly enriched than those targeted by one sgRNA. These data indicate the high efficiency and specificity of CATE.

**Figure 3.**
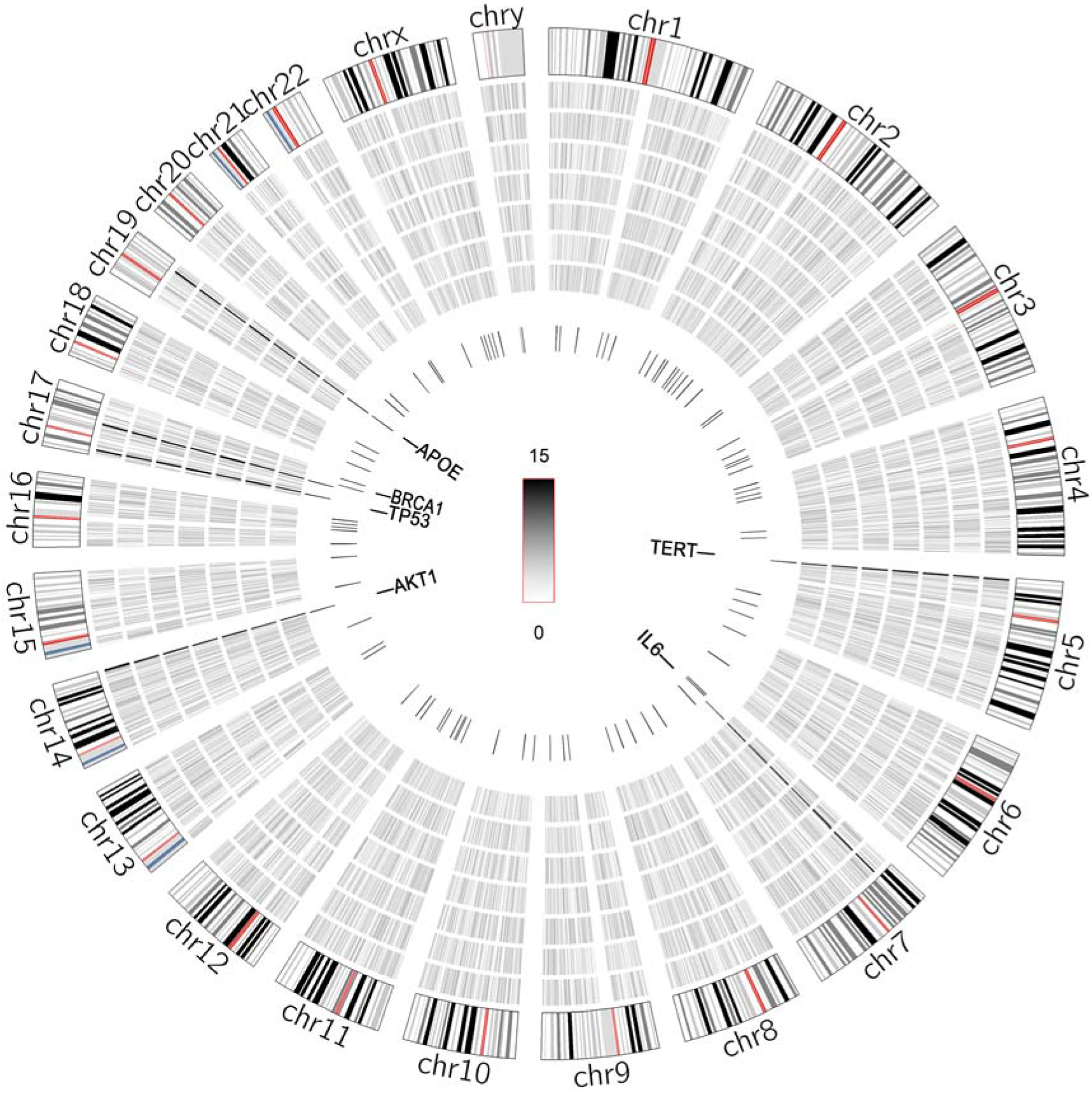
Distribution of mappable reads of seven DNA samples in human genome. From outer to inner layers, chromosome map, the CATE-seq reads density of 293T, 293Tm, HepG2, HL7702, HeLa, SiHa, and C-33A DNA samples, respectively, sgRNA targets, and the predicted off-targets of sgRNAs. The reads density refers to the reads numbers in each 1-Mb window. The log2 value of reads density was then calculated and shown as Circos. The vertical lines in the most inner layer are positions of the predicted off-targets of sgRNAs. It seems that a predicted off-target overlaps with the APOE gene in this figure. In fact, it is distant from the APOE gene locus as far as 1,140,004 bp (off-target position is chr19:44270841 but APOE gene position is chr19:45410845).

**Figure 4.**
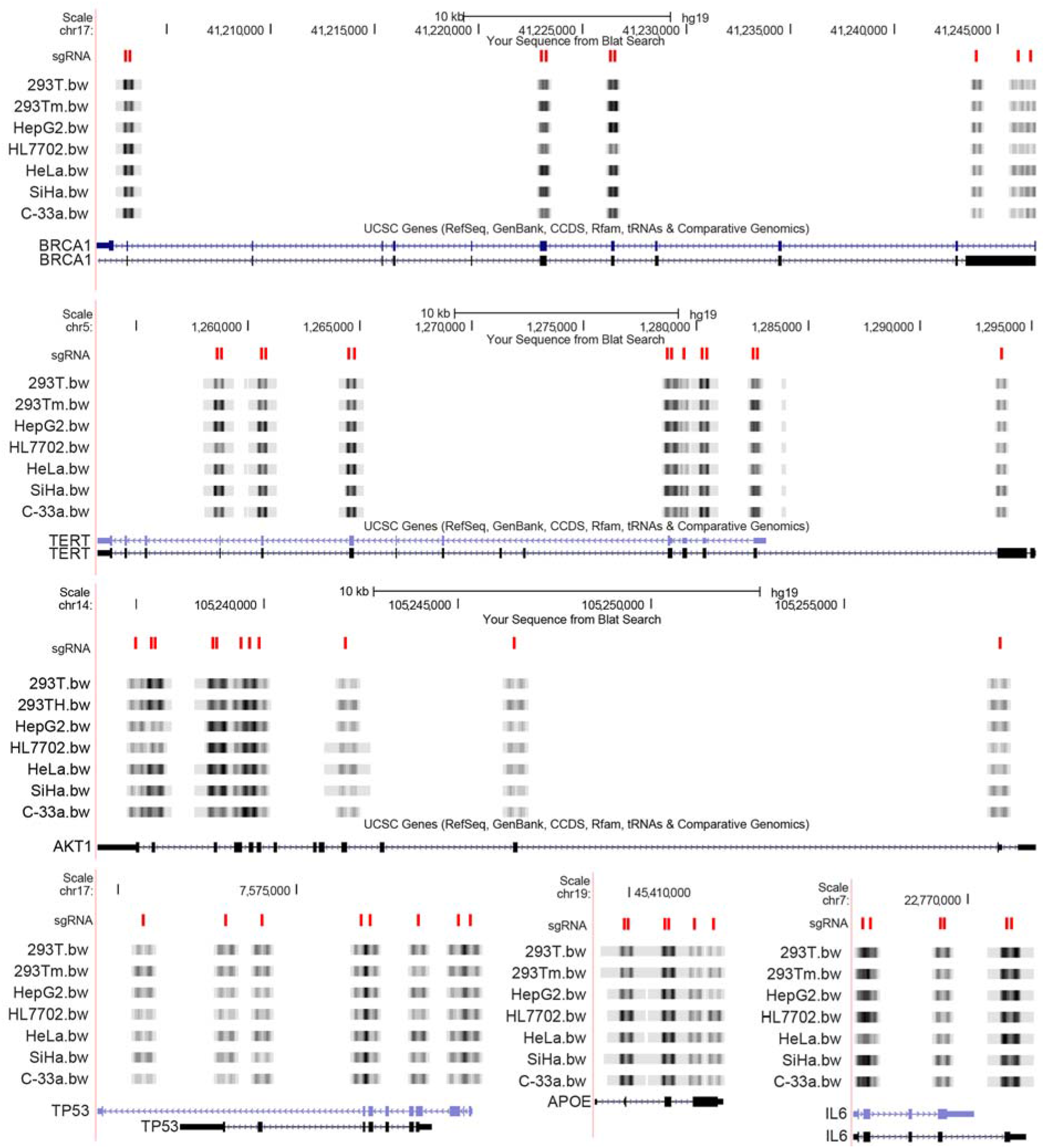
Distribution of CATE-seq reads at the loci of target genes. Based on the mapping results, the reads data in BigWig files were displayed with the UCSC Genome Browser. The BigWig files were used as customized Track files and hg19 was used as the reference genome. The mappable reads distribution in the whole 6 target gene loci in 7 gDNA samples of 6 cell lines were shown. The gray scale indicates the reads density. The whole gene loci and all exons of 6 target genes were shown. The positions of all csgRNA targets were shown as vertical lines in red in the track of sgRNA.

### Reads coverage to the targeted exons

The distribution of all mappable reads in 6 genes in 7 DNA samples were visualized with the UCSC genome browser to find the reads coverage to the targeted exons. The results indicated that the CATE-seq reads were highly enriched at the position of sgRNA targets (Figure 5; Figure S6-S10). The exons targeted by two sgRNAs showed higher enrichment than those targeted by one sgRNA. The full length of target exons was highly covered by the mappable reads. Additionally, many intronic sequences flanking the target exons were also highly enriched and sequenced. In this study, a single sgRNA was designed for as many as 17 exons (48.5%). It was found that both the full length of target exons and the partial sequence of flanking introns were highly covered by the CATE-seq reads enriched by these single sgRNAs (Figure 5; Figure S6-S10). The length of these exons is 85–246 bp (Table S9); however, the sequences covered by over 50000 reads were 495-582 bp in length (Table S9). For other 18 exons, 17 exons were designed with two sgRNAs and only one exon was designed with three sgRNAs (Table S9). The length of these exons is 94–310 bp (Table S9); however, the sequences covered by over 50000 reads were 495–865 bp in length (Table S9). These data suggest that one sgRNA is sufficient for targeted enrichment of most exons in human genome. It is not necessary to design two sgRNAs for 89% of exons that were designed with two sgRNAs in this study (Table S9). Two or more sgRNAs should be designed only for a few long exons (Table S9). These data also suggest that sgRNA can be designed in introns flanking target exons that are too short to design sgRNA.

**Figure 5.**
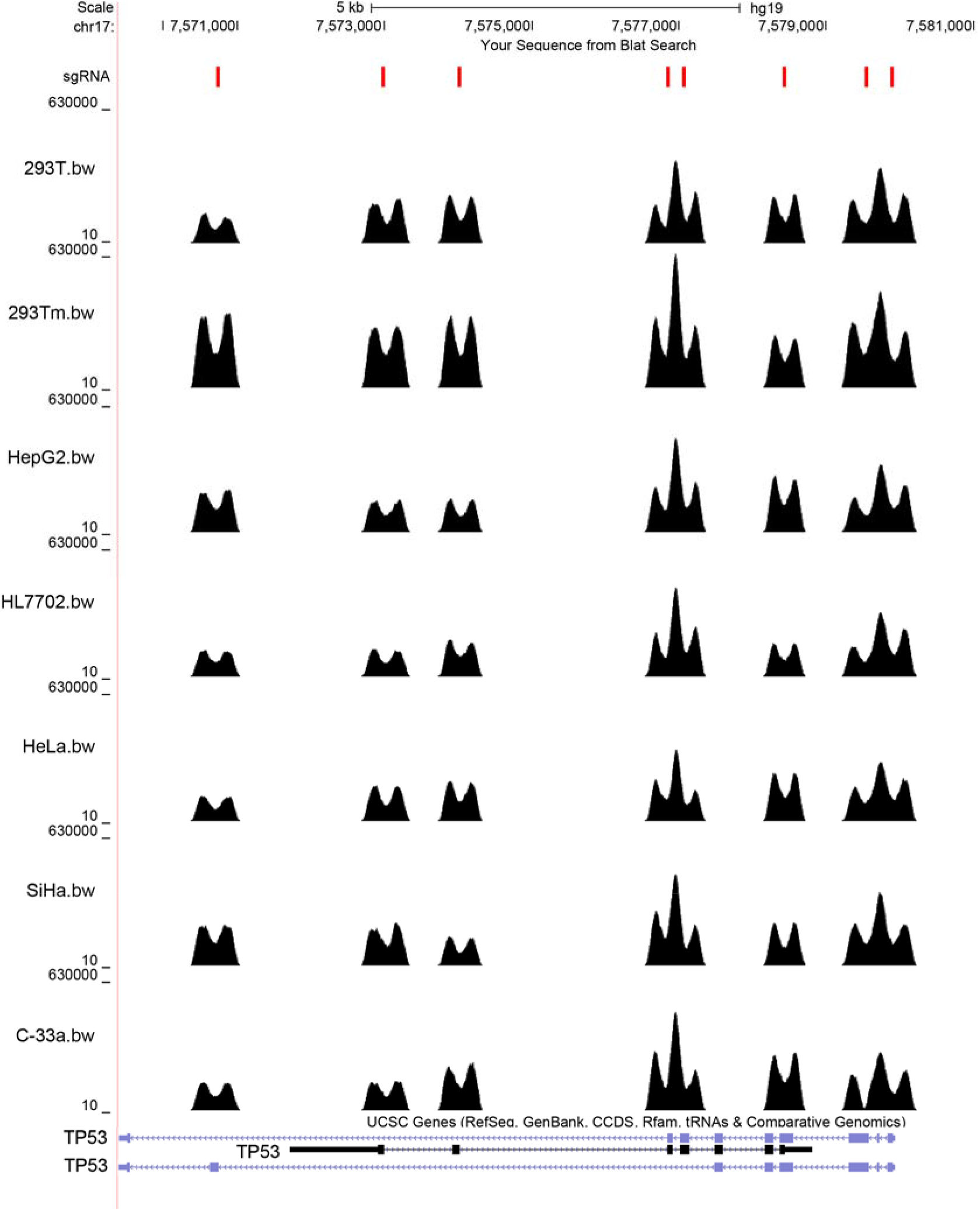
Reads distribution in TP53 gene locus. Reads distribution of 7 DNA samples were shown. The positions of all csgRNA targets (vertical lines in red), the whole gene regions and all exons of 6 target genes were shown.

The reads distribution in the full length of all target exons of 6 genes in each DNA sample was further checked at a single-base resolution. The results demonstrate that the full length of exons are highly covered by large numbers of reads (Figure 6). The comparison of reads distribution in 6 genes in 6 DNA samples revealed that the enrichment efficiency of dCas9-csgRNA in variant DNA samples were highly stable (Figure 6). These data suggest that CATE is applicable to targeted sequencing for finding somatic mutations.

**Figure 6.**
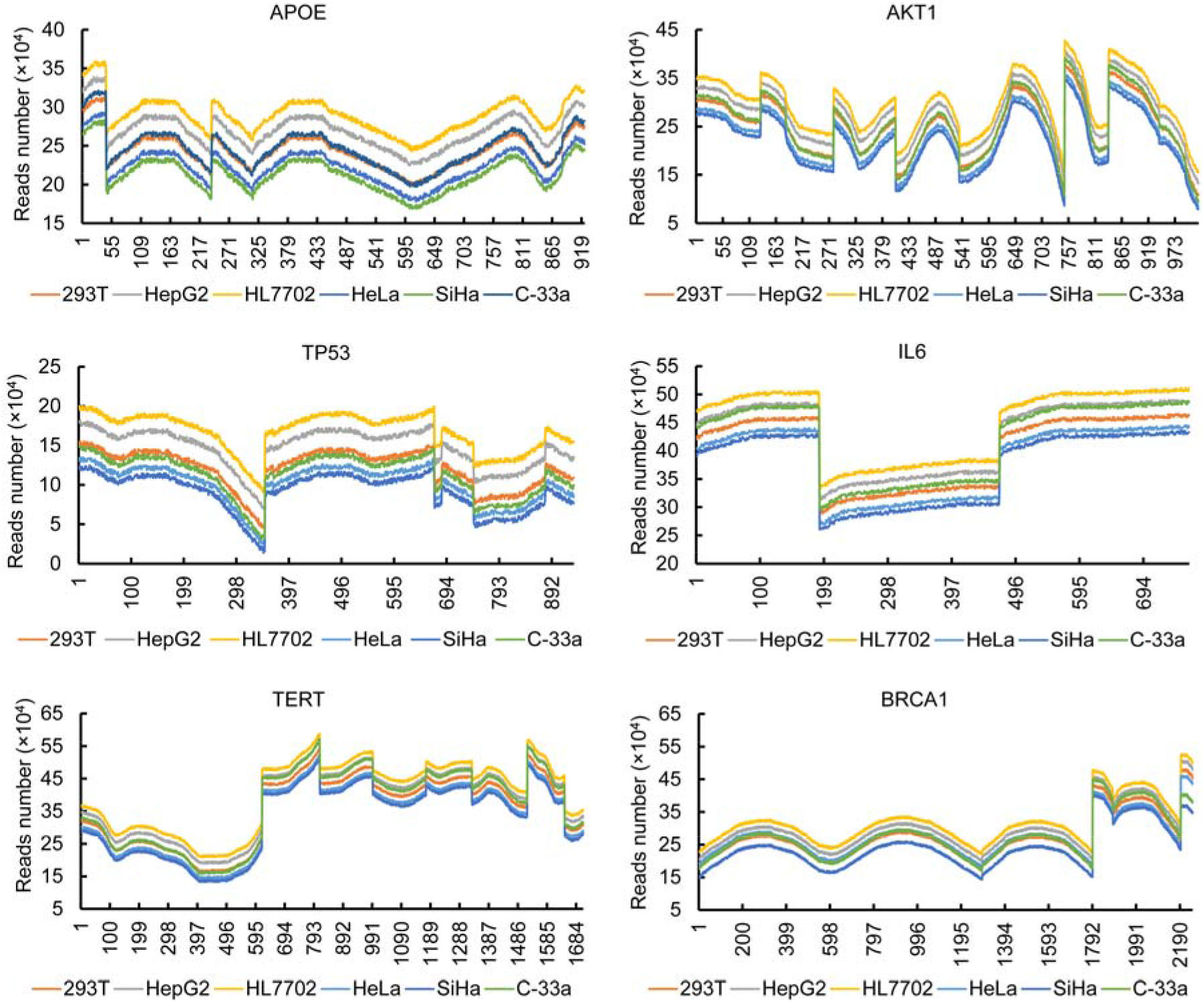
Reads distribution in the targeted exons of 6 genes in 6 cell lines. Reads distribution in each target exons of 6 genes in 6 cell lines were counted at a single-base resolution. Only the targeted exons were shown. The unit of abscissa is base pair (bp), indicating the full length of spliced target exons.

### Statistic of mutations occupied by over 50000 reads

Mutation analysis was next performed with CATE-seq data for finding the potential somatic mutations in the target exons. The results revealed that most mutations exist in introns flanking the sgRNA-targeted exons. Only a few mutations were found in in target exons (Table S10). It was found that HepG2, C-33a, and HeLa cells contained the rs1042522 mutation (Figure 7A). This SNP locates in the coding region of TP53 gene and is a known high-risk mutation associated with multiple tumorigenesis (40,41). This mutation exists in hepatoma cell HepG2 but not in normal liver cell HL7702. In three cervical cancer cells, HeLa is a HPV18-positive cell (42), SiHa is a HPV16-positive cell (43), and C-33a cell is a HPV-negative cell (44). According to the CATE-seq results and the HPV infection, it can be reasoned that the HeLa cell carcinogenesis may be caused by a combination of the HPV infection and TP53 mutation, the SiHa cell carcinogenesis may be caused by the HPV infection alone, but the C-33a cell carcinogenesis may be caused by the TP53 mutation.

**Figure 7.**
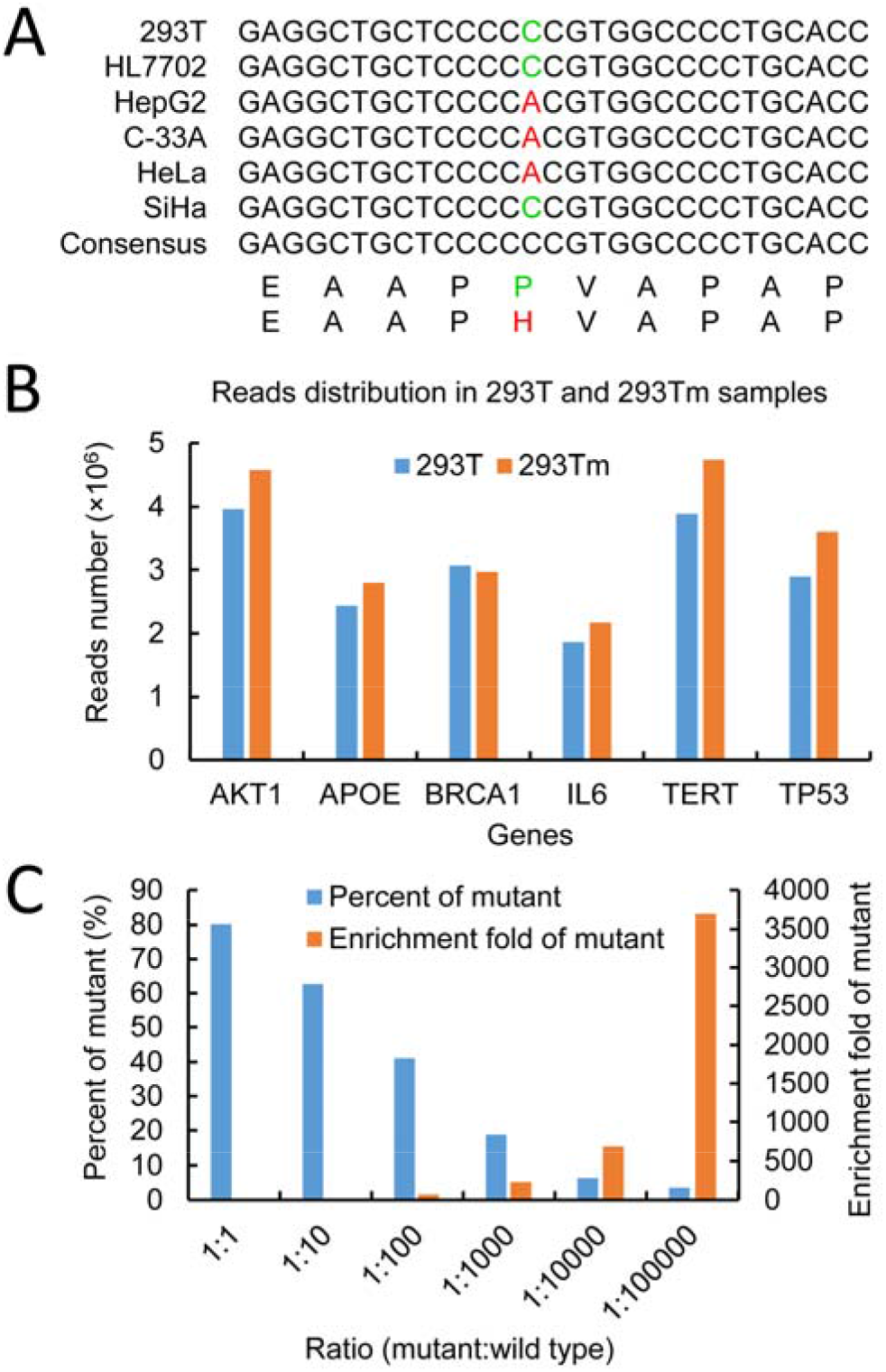
CATE performance in mutation identification, throughput and sensitivity. (A) A mutated site found by CATE-seq in this study. This is a hotspot mutation widely reported in many cancers. (B) Reads distribution in the enriched 293T and 293Tm gDNA samples. The reads mapped to each target gene in the enriched 293T and 293Tm gDNA samples were counted and shown. (C) Targeted enrichment of mutant TERT promoter sequence. The mutant and wild-type TERT promoter sequences were mixed at the different ratios. The mutant TERT promoter sequence was then enriched from these mixtures with CATE. The enriched DNA was analyzed by ARMS-qPCR. The percentage and enrichment fold of mutant TERT promoter sequence in the enriched DNA were calculated.

### Enhancing CATE throughput with more complex csgRNA library

In the enrichment of 293T DNA sample, 54 csgRNAs were divided into 4 groups. Each group was used for an independent targeted enrichment of gDNA. Then, the enriched gDNAs were pooled together and used as the final CATE product of 293T cell. However, whether more complex csgRNA pool could be used to enrich many targets simultaneously is important to simplifying operation and enhancing throughput of CATE method. In the enrichment of 293Tm DNA sample, all 54 csgRNAs were pooled together and used to enrich 293Tm gDNA. The reads distributions in 6 genes in the 293Tm and 293T DNA samples were compared. As a result, no significant difference was found in two ways of enrichment, indicating that inclusion of more different csgRNAs in a CATE reaction did not affect targeted enrichment. That is, there was no mutual interference among csgRNAs. This is helpful for enhancing the throughput of CATE-seq (Figure 7B).

### Further validation of specificity and sensitivity of CATE

For further evaluating its specificity, CATE was used to enrich a mutant TERT promoter from an equimolar mixture of a wild-type (TERT-P) and a mutant (TERT-P-mut) TERT promoter fragment. The enriched DNA was analyzed by PCR amplification and the followed cloning sequencing. The results revealed that there were 19 TERT-P-mut and 1 TERT-P clones in 20 sequenced positive clones (see supplementary sequence), indicating that the mutant sequence was highly and specifically enriched by CATE. The mutant TERT promoter was also enriched from a series of mixtures of the TERT-P-mut and TERT-P at different ratios. The enriched DNA was quantitatively analyzed with ARMS-qPCR. The results showed that TERT-P-mut was enriched from all samples (Figure 7C). Especially, TERT-P-mut diluted in TERT-P at a lowest ratio (1:1,000,000) was still successfully enriched, with an enrichment fold up to 3691 (Figure 7C).

### Verification of CATE-seq by large-scale targeted enrichments

For verifying the scalability of CATE-seq, a second and third CATE-seq assays were then performed. In a second CATE-seq assay, 367 sgRNA targeting 339 exons of 186 genes were designed. These genes are closely related with carcinogenesis and collected by COSMIC. The individually-prepared 367 csgRNAs were mixed together at the same micrograms to associate with dCas9. The formed dCas9-csgRNA complex pool was used to enrich their targets from the tagmented gDNAs of the same six cell lines. The NGS revealed that all targets were highly enriched from all gDNA samples (Table S11–S13; Figure 8), indicating the good scalability of our method. For further simplifying the CATE-seq assay, 367 csgRNAs were then prepared in a high-throughput format, in which all F2 and F3 primers were respectively mixed at the same mole and used as F2 and F3 primers in the second and third round of PCR. A pool of 367 csgRNA transcription templates was thus prepared, which was then used to transcribe 367 csgRNAs together and form a csgRNA pool. This high-throughput protocol for preparing large numbers of different csgRNAs is highly time- and labor-effective. The targets in the tagmented gDNAs of two cell lines (293T and HL7702) were then enriched with this new csgRNA pool. The results indicated that all targets were enriched (Table S11-S13; Figure 8); however, variant targets were enriched in different efficiency (Table S13). This may be caused by the different amplification efficiency of variant csgRNA templates in the three-round PCR reaction, which was supported by the almost identical enrichment efficiency of variant targets from two gDNA samples (293Tp367m and HL7702p367m in supplementary Table S13). This limitation may be improved by including less F2 and F3 primers in a PCR reaction (such as 100). It was found that 35 target exons of 6 genes in 6 gDNA samples were similarly enriched in the first and second CATE-seq assays that used a csgRNA pool of 54 and 367 csgRNAs, respectively (Figure 3 and Figure S11), indicating the good reproducibility of CATE-seq method. The mutations in the enriched exons and flanking introns could be identified using the CATE-seq data (Table S14).

**Figure 8.**
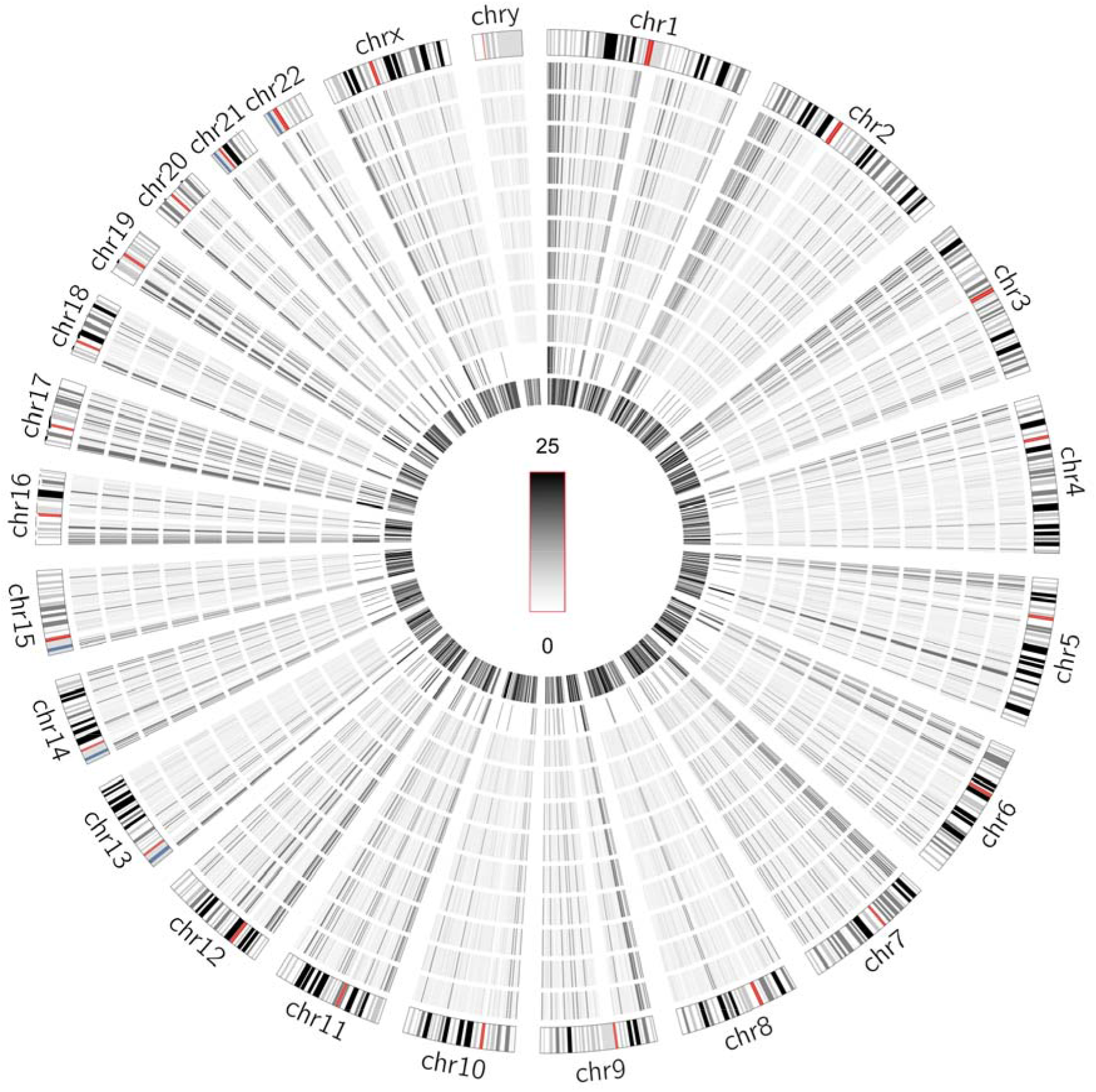
Distribution of mappable reads of nine DNA samples in human genome. 339 exons of 186 genes were enriched with 367 sgRNAs. From outer to inner layers, chromosome map, the CATE-seq reads density of 293T, HepG2, HL7702, HeLa, SiHa, C-33A, 293T50m, 293Tp367m, and HL7702p367m DNA samples, respectively, sgRNA targets, and the predicted off-targets of sgRNAs. The reads density refers to the reads numbers in each 1-Mb window. The log2 value of reads density was then calculated and showed as Circos. It seems that there are some predicted off-targets overlapped with enriched positions in this figure. In fact, they are distant from sgRNA targets.

In a third CATE-seq assay, a total of 2302 csgRNAs targeting 2031 exons of 451 protein-coding genes were designed (Table S3). These genes are collected in a current FDA-approved NGS-based largest cancer gene detection panel, MSK-IMPACT™, which consist of cancer-related 468 genes. MSK-IMPACT is a custom hybridization capture-based assay encompassing all genes that are druggable by approved therapies or are targets of experimental therapies being investigated in clinical trials at Memorial Sloan Kettering Cancer Center (MSKCC), as well as frequently mutated genes in human cancer (somatic and germline mutations) (45–48). Importantly, csgRNAs were only designed for those exons containing mutations with close relationship with carcinogenesis and collected by COSMIC. The linear *in vitro* transcription templates of 2302 csgRNAs were individually prepared in 24 96-well PCR plates by PCR amplification using primers listed in Table S4. CgRNAs were then prepared in two 96-well PCR plates by *in vitro* transcription using 20 templates per reaction. A total of 2302 csgRNAs were finally mixed into 7 pools of about 350 csgRNAs to associate with dCas9, respectively. The formed dCas9-csgRNA complex pools were used to enrich their targets from the tagmented gDNAs of two cell lines (HepG2 and HeLa). The NGS revealed that all targets were highly enriched from two gDNA samples (Table S15 and S16; Figure 9), indicating the high scalability of CATE-seq method. Finally, the mutations in the enriched exons and flanking introns were identified using the CATE-seq data (Table S17).

**Figure 9.**
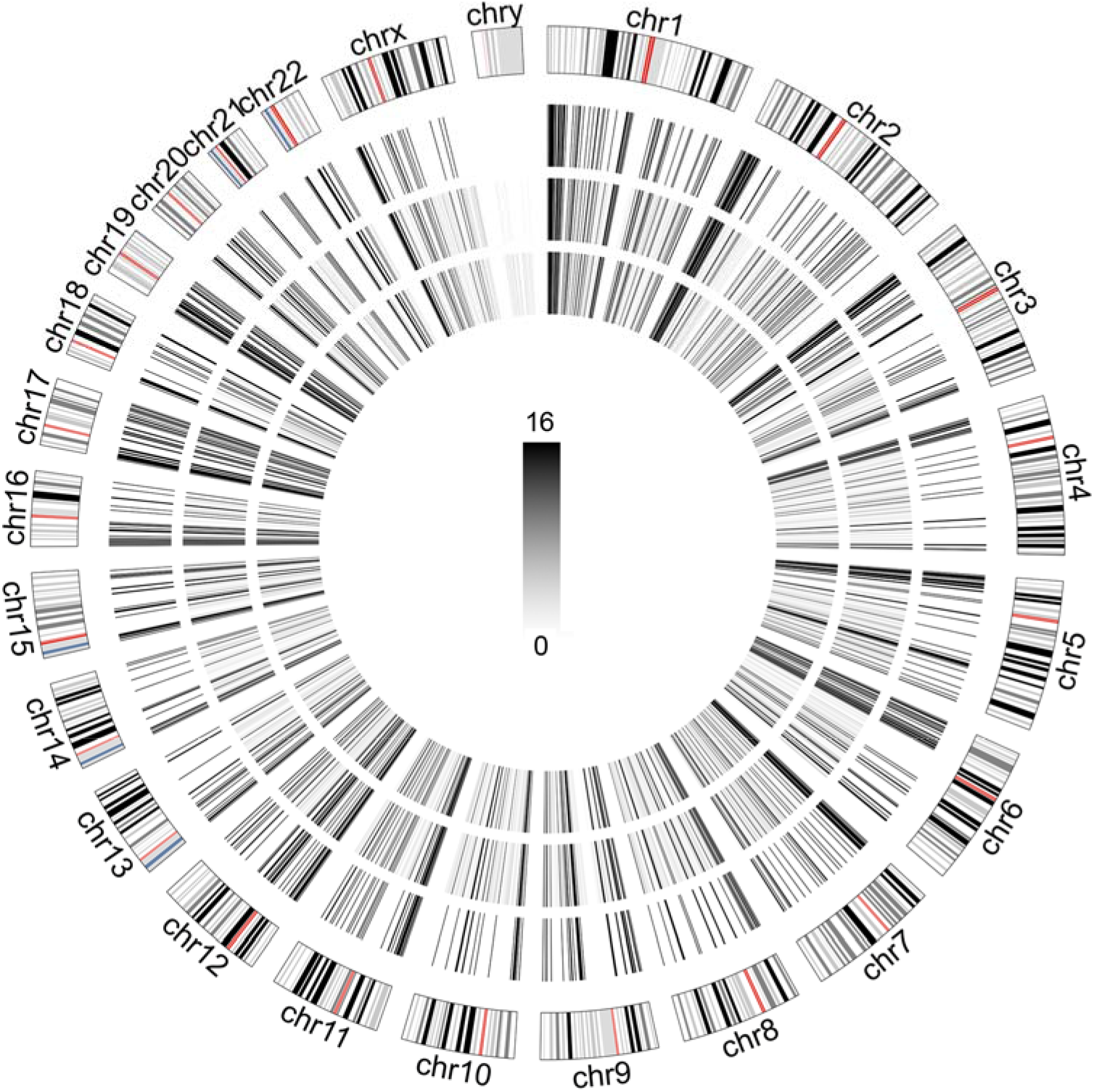
Distribution of mappable reads of two DNA samples in human genome. 2031 exons of 451 genes were enriched with 2302 csgRNAs. From outer to inner layers, chromosome map, sgRNA targets, HepG2 reads density, and HeLa reads density. The reads density refers to the reads numbers in each 1-Mb window. The log2 value of reads density was then calculated and showed as Circos.

## DISCUSSION

With the rapid development of CRISPR technology, besides its wide applications in gene editing and regulation, it has been rapidly applied to targeted sequencing in the past two years. Several CRISPR-based methods for targeted sequencing have been developed, including STR-Seq (49), CRISPR-DS (50), and CRISPR-Cap (51). Especially, specific megabase segments of genomic DNA can be enriched by CRISPR-mediated isolation (52). However, these methods used Cas9-sgRNA to cut out target DNA fragments from gDNAs. The excised small or large target fragments were then isolated by hybridizing with probes coupled in flowcell (49), or performing size selection using SPRI beads (50) or pulse field gel electrophoresis (PFGE) (52). Only in the CRISPR-Cap method, the excised target DNA fragments were isolated with the biotinylated sgRNAs using streptavidin-coupled magnetic beads (51). However, biotin molecules incorporated in sgRNAs as biotinylated nucleotides in *in vitro* transcription may interfere the Cas9-sgRNA association and digestion to its targets. Additionally, different labeling efficiency of variant sgRNAs may affect their capturing efficiency.

This study developed a new CRISPR-based targeted sequencing method, CATE-seq. Compared with the current CRISPR-based methods, CATE-seq has several unique features. CATE-seq did not use Cas9 to cut gDNA as other methods. CATE-seq used dCas9 and a simply engineered label-free sgRNA (csgRNA) to enrich target DNA fragments from any gDNA libraries sheared by all methods such as tagmentation, sonication and enzymatic digestion. Although a 6×His tag-fused dCas9 protein was recently used to isolate target cell-free DNA (cfDNA) by depending on anti-6×His-coupled magnetic beads (53), the lower binding affinity between 6×His tag and antibody is unbeneficial to high specific and stable capture. In contrast, CATE-seq used the high-affinity interaction between biotin and streptavidin to capture dsDNA-dCas9-csgRNA complex, which obtained high percentage of targeted reads (nearly 90%).

CATE has significant advantages over the hybridization-based targeted enrichment (HBTE) methods. First, CATE has high simplicity. Different from the HBTE strategies, CATE is a targeted enrichment strategy free of traditional hybridization. All HBTEs start with a denaturation of DNA sample at high temperature (95°C) and undergo a long-time single- or two-round hybridization process (1.5–4 h or 16–72 h) at 47– 65°C(54). After hybridization, the captured sample has to be washed several times (e.g. twice) at high temperature (e.g.47°C) and several more times (e.g. three times) at RT using the manufacturer’s buffers (54). However, CATE needs no denaturation and hybridization of DNA sample. CATE provides an easy targeted enrichment strategy, which can be rapidly finished in as few as 2 h. The whole CATE procedure can be operated at RT without requiring any HBTE-needed hardware (e.g. hybridization station). CATE also needs no any HBTE-needed high-cost DNA chips or lengthy biotin-labeled capture probes. Another advantage of CATE is that CATE directly captures dsDNA but HBTE captures ssDNA. This means that one sgRNA can capture two strands of target DNA; however, two different sets of oligo capture probes have to be designed to capture two strands of target DNA. CATE therefore greatly simplified the complexity of capture probe (sgRNA) design and selection.

A key feature of CATE is that it captures dsDNA-dCas9-csgRNA complexes on magnetic beads by a rapid RNA:DNA annealing at RT. This facile process is dependent on the advantage of better annealing efficiency and stability of RNA:DNA hybrids than those of DNA:DNA hybrids (55). It should be noted that the magnetic isolation step in CATE was performed at RT. It was found that the dsDNA-dCas9-csgRNA complex could be captured by magnetic beads in high efficiency, suggesting the high-efficiency annealing between csgRNA CS and beads-coupled CP at RT. This is the most advantageous step of CATE. It should be mentioned that the short csgRNA CS (24 bp) used in this study was an artificially designed sequence, which has no homology with any human genomic DNA sequences. This can prevent potential interference from possible single-stranded gDNAs and contaminated RNAs. This study revealed that the 24-bp csgRNA CS functioned excellently in CATE.

SgRNA is the main experimental material for CATE. Perfect sgRNA designing is critical to successful CATE-seq. In sgRNA designing, the reported somatic mutations in target exons were carefully checked. Many sgRNAs were designed but those with targets without mutations were selected, which prevents the potential interference of mutations to CATE. However, the mutation-specific sgRNA can be designed for enriching those known disease-causing mutations, as mutant TERT promoter was enriched in this study. Due to extensive presence of PAM sequences in human genome, sgRNA design and selection are less of an issue for CATE. For preparing csgRNA simply and rapidly by *in vitro* transcription, a new sgRNA transcription template preparation method recently developed by our lab was used (56). The sgRNA transcription template can be rapidly prepared by a three-round PCR protocol. Using this protocol and *in vitro* transcription, many csgRNAs can be prepared in two days. More importantly, a multitude of sgRNAs can be prepared in library using the procedure, which greatly simplifies the sgRNA preparation process.

CATE has high specificity. The high specificity of CATE method is dependent on high specific interaction between Cas9-sgRNA and its target DNA, which greatly differs from the current HBTE sequencing methods. This avoid nonspecific enrichment resulted from nonspecific hybridization between capture probes and the denatured DNA sample. It is difficult to design a multitude of high specific capture probes to various targets in HBTE methods, because the probes have to have the similar annealing temperature for obtaining similar hybridization efficiency. On chip, the nonspecific annealing between capture probes and DNA sample may result in high nonspecific enrichment and noise signal. In solution, besides the nonspecific annealing between capture probes and DNA sample, the nonspecific annealing among capture probes may further reduce enrichment efficiency. Additionally, sequences with high A-T or G-C content can be lost by HBTE through poor annealing and secondary structure formation (15). The potential bias for repetitive elements can also cause more non-uniquely mapped reads. The high specificity of CATE was fully verified by the results of this study. In total, nearly 90% of mappable reads were sgRNA targets. Additionally, the mutant TERT promoter sequence highly diluted in wild-type TERT promoter sequence (1:1,000,000) was specifically enriched by CATE. Importantly, there is only one base difference between the wild-type and mutant TERT promoter sequences, indicating the high specificity of Cas9-sgRNA binding with its target DNAs. These data suggest that the gDNA fragments with rare and low-frequency disease-related SNPs can be specifically isolated from the detected DNA samples by CATE. For example, the mutant TERT promoter sequence used in this study reactivates the expression of telomerase in most of cancers (57,58). Therefore, CATE provides a potential tool for clinical diagnosis, especially for noninvasive prenatal testing (NIPT) and liquid biopsy.

CATE has high sensitivity. This study demonstrated the high sensitivity of CATE by a gradient enrichment assay. A mutant TERT promoter sequence was mixed into the wild-type TERT promoter sequence at the different ratios, with the lowest dilution of 1:1,000,000. It was found that the mutant TERT promoter sequence could be successfully enriched from the lowest dilution, with the enrichment fold of 3691 times. As a representative HBTE, it was reported that CAPP-Seq could detect one mutant DNA molecule from in 10,000 healthy DNA molecules (5). Due to its high sensitivity, CATE may be used to detect rare mutations such as those in cell-free fetal DNA (cffDNA) and cell-free tumor DNA (ctDNA) in liquid biopsy. The high sensitivity of CATE method was also revealed by the used low-amount input DNA. As little as about 5 ng of gDNA was used in all enrichments. In contrast, 10–15 μg of starting DNA material was required by array-based HBTE to drive the hybridization to completion (6), whereas 50 ng to 100 ng of good-quality input DNA was needed by most of commonly used in-solution HBTEs including SureSelect (Agilent), Nextera (Illumina), TruSeq (Illumina), and SeqCap EZ (Roche Nimblegen) (9). Even in the newest in-solution HBTE, the SureSelect Human All Exon V7 (Agilent), 10 ng of input DNA is still needed. Requirement of more input DNA challenges the applications of HBTEs to some important clinical samples such as formaldehyde-fixed paraffin-embedded (FFPE) tissue.

CATE has high throughput. First, many different sgRNAs can be prepared in a library without mutual interference in enrichment, which enhances the sgRNA preparation throughput. Secondly, many targets in a DNA sample can be captured by a csgRNA cocktail in parallel. By comparing the CATE-seq results obtained with csgRNA pools containing variant numbers of different csgRNAs (50 and 367), it was found that the complicated csgRNA cocktail could be used to enrich various targets in a high-throughput format without mutual interference. Thirdly, the barcoded DNA samples can be mixed together and enriched by CATE as a single DNA sample, which can greatly simplify operation, enhance throughput, and reduce bias.

The gDNA sheared by Tn5 transposome (a process now named as tagmentation) was used as input DNA sample in this study; however, all DNA fragments sheared by any other methods can be used in CATE-seq (Figure S12), such as those produced by endonuclease digestion and sonication. Additionally, the naturally degraded DNA can also be used by CATE-seq, such as blood cfDNA and FFPE DNA. When applied to cfDNA, CATE-seq is beneficial to identify disease-causing mutations in liquid biopsy. In this study, the NGS library construction process of CATE-seq was designed to preferentially use our recently developed SALP method (36). This is a single-stranded library construction method, which is qualified to construct NGS library of any sources of DNA samples, especially high-degraded DNA such as cfDNA. Therefore, CATE-seq may be beneficial to analyze cfDNA in the future liquid biopsy.

This study revealed that CATE-seq can also be used as a new method to characterize the targeting of Cas9-sgRNA. In this study, the genomic mapping of all CATE-seq reads revealed that nearly 90% of mappable reads were sgRNA targets. The remained mappable reads were originally suspected as off-targets. However, these reads were almost evenly and randomly distributed in the whole genome (Figure 3, 8 and 9). Their genomic distribution had no correlation with the predicted off-targets (Figure 3 and 8). Therefore, these reads was reasoned coming from the little non-specific adsorption of input DNA to magnetic beads, which may be overcome by more sufficient blocking and stringent washing in CATE. These results indicate that the dCas9-csgRNA has high target specificity, which ensures the high specificity of CATE.

In summary, this study developed a new technique of CRISPR-based targeted sequencing, CATE-seq. A new kind of sgRNA (csgRNA) with a short CS that can anneal with a CP immobilized on magnetic beads was engineered. The gDNA library were firstly incubated with the pre-assembled dCas9-csgRNA complex, allowing target dsDNA fragments to be specifically bound by the dCas9-csgRNA complex. The dsDNA-dCas9-csgRNA complexes were then isolated with the CP-coupled magnetic beads. By using the technique, three different scales of enrichments were successfully performed. In a largest-scale enrichment, as many as 2031 target exons of 451 genes were specifically enriched from two different gDNA samples with 2302 variant csgRNAs. This study verified the high simplicity, specificity, sensitivity, throughput, and scalability of this technique. This study thus provides a new powerful tool for targeted sequencing that has significant advantages over the current hybridization- and CRISPR-based methods.

## DATA AVAILABILITY

The reads data of CATE-seq are available at NCBI GEO with the accession number: GSE119994.

## SUPPLEMENTARY DATA

Supplementary Data are available online. Supplementary data include three supplementary methods, fourteen supplementary tables, twelve supplementary figures, and two cloning sequencing results.

## ACKNOWLEDGMENTS

We thank two anonymous reviewers for their kind comments and suggestions to manuscript revision.

## Authors contributions

Xinhui Xu designed sgRNA and performed targeted enrichment and data analysis. Qiang Xia, Shuyan Zhang, and Jinliang Gao prepared csgRNA. Wei Dai cultured cells and prepared transposome and tagmented genomic DNA. Jian Wu performed data analysis. Jinke Wang conceptualized the project, designed the research and wrote the manuscript. All authors read and agreed the content of the final manuscript.

